# Identification of distinct rodent-associated adenovirus lineages from a mixed-use landscape in the Western Ghats biodiversity hotspot

**DOI:** 10.1101/2024.05.18.594665

**Authors:** B.R. Ansil, Avirup Sanyal, Darshan Sreenivas, Kritika M. Garg, Uma Ramakrishnan, Balaji Chattopadhyay

**Author notes:** These first authors contributed equally to this manuscript (B.R. Ansil, Avirup Sanyal).

## Abstract

Shifts in land-use patterns and increased human-livestock-wildlife interactions have generated numerous possibilities for pathogen spillover. This demands increased efforts of pathogen surveillance in wildlife, especially in changing landscapes with high biodiversity. We investigated adenovirus diversity in small mammals, an understudied host taxon, from a forest-plantation mosaic in the Western Ghats biodiversity hotspot. Using PCR-based screening followed by Sanger sequencing and phylogenetic analyses, we attempted to detect and characterize adenovirus diversity in seven species of small mammals. We observed high prevalence (up to 38.8%) and identified five lineages of adenoviruses with unique mutations in the endemic and dominant small mammal species, *Rattus satarae*. These lineages significantly differed from other known Murine adenoviruses (p-distance > 25%), indicating the likelihood of novel adenovirus diversity in this endemic small mammal. Collectively, our results highlight the potential for unexplored diversity of DNA viruses like adenovirus in poorly explored host taxa inhabiting human-used landscapes and its zoonotic implications.

## 1. Introduction

Adenoviruses are a diverse group of non-enveloped, double-stranded DNA viruses known to infect vertebrates, including humans (MacNeil et al., 2023). These generalists are speculated as efficient pathogens of global threat causing mild to severe infections, ranging from gastrointestinal diseases, common cold, encephalitis, respiratory and hemorrhagic diseases often resulting in death (Saint-Pierre Contreras et al., 2023). Research efforts have mainly focused on understanding adenovirus infections in humans and livestock, (e.g. Ghebremedhin, 2014), while their prevalence and diversity in animal reservoirs, specifically in small mammals remain relatively unexplored. Existing eco-epidemiology of adenoviruses in small mammals (Ochola et al., 2022; Zheng et al., 2016) provides poor insights into the diversity of adenoviruses in species-rich mammals, specifically in India.

As the most species-rich mammalian group, small mammals are predicted reservoirs for many novel zoonotic pathogens (Luis et al., 2015). Anthropogenic disturbance can increase small mammal densities, enhancing the risk of emerging infectious diseases (Mescht et al., 2013). Regions where high mammal diversity intersects with land-use change can result in novel species assemblages, promoting subsequent zoonotic spillovers (Allen et al., 2017). Therefore, understanding the diversity of generalist viruses like adenovirus in small mammal communities is of significant public health interest. Here, we investigated the prevalence and diversity of adenovirus in a small mammal community, comprising rodents and shrews in a forest-plantation mosaic located in the biodiverse Western Ghats in southern India, which is also inhabited by other free-ranging mammals, livestock, and humans. Due to their proximity to humans and potential interactions with other wildlife species, in such mixed-use landscapes, small mammals have the prospect of acting as natural maintenance and intermediate hosts for adenovirus transmission. By investigating adenovirus in these often-overlooked hosts, we aimed to gain valuable insights into the broader epidemiologic implications of small mammals in changing landscapes.

## 2. Methods

### 2.1 Study area and samples

We leveraged already collected small mammal (rodents and shrews) samples from Kadamane forest-plantation mosaic in Karnataka state in Southern India (Ansil et al., 2021). This region is part of the Western Ghats biodiversity hotspot, has high human density and rapid modification of natural habitats and high human-animal interactions. This makes it an important landscape to understand viral diversity associated with wildlife. Small mammals were captured from different land-use types; forest fragments, grasslands, and human habitations (Figure 1) between January and March 2018 (NCBS-Institutional Animal Ethics Committee approval-NCBS-IAEC-2016/10-[M]). Samples included 136 small mammal individuals representing seven distinct species, (previously reported in Ansil et al., 2021, 2023); *Rattus satarae, R. rattus, Mus cf. fernandoni, M. cf. famulus, M. cf. terricolor, Platacanthomys lasiurus*, and *Suncus niger*. In the forests, *R. satarae* was the most abundant species (n= 67, Figure 1), followed by *M. cf. fernandoni* (n=2) and *M. cf. famulus* (n=2). *P. lasiurus* was rare, and only one individual was captured during the sampling. In grasslands, *M. cf. fernandoni* (n=21) and *M. cf. famulus* (n=20) exhibited nearly equal abundance, whereas *M. cf. terricolor* had relatively lower abundance (n=6). In human habitation, *R. rattus* was most abundant (n=10), followed by *M. cf. famulus* (n=5) and *S. niger* (n=2).

**Figure 1:**
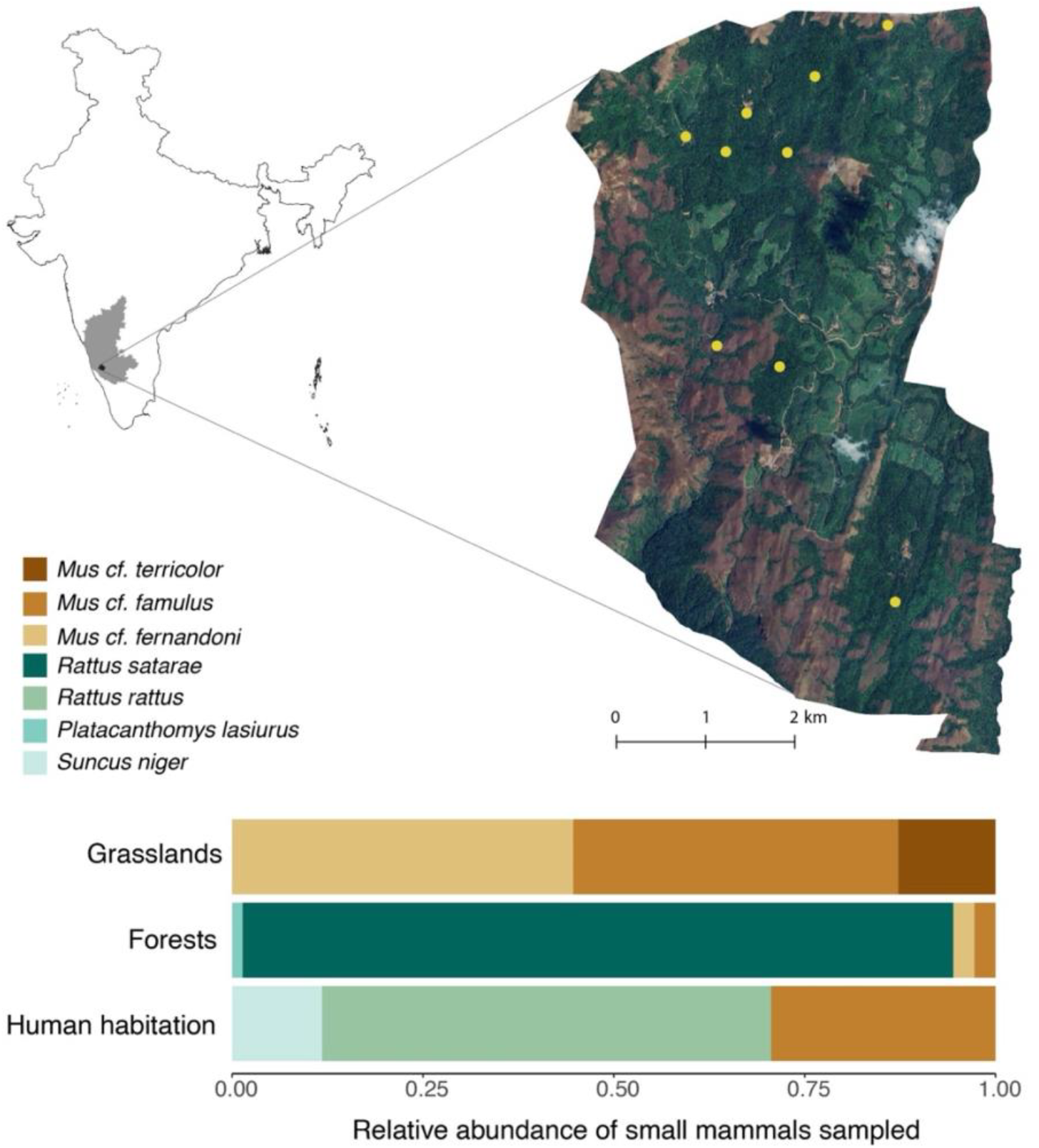
Study area in the Western Ghats, in Southern India. The gray shaded area and enlarged area with satellite imagery shows Karnataka state and Kadamane forest-plantation mosaic respectively. The stacked bar plots show relative abundance of various small mammal species captured during the sampling. Except for *Platacanthomys lasiurus*, all the small mammal samples were tested for adenovirus.

### 2.2 Adenovirus screening

We extracted DNA from pooled tissues (liver, spleen, lungs, kidney, and intestine), oral swabs, and rectal swabs, using Quick-DNA/RNA Miniprep Plus Kit-D7003 (Zymo Research Corporation) following manufacturer’s protocol. All the DNA samples were screened for the presence of adenovirus using a set of degenerate primers targeting a short fragment of 270 base pair (bp) of the DNA polymerase (DPOL) gene of the adenovirus (Wellehan et al., 2004). We purified the prospective positive samples, sequenced them at the NCBS Sanger sequencing facility. The study was approved by NCBS Institutional Biosafety Committee (TFR: NCB:23_IBSC/2017).

Chromatograms of Sanger sequencing reads (DPOL gene-forward and reverse) for each positive sample were visually inspected, primer binding sites were trimmed, and a consensus sequence was generated in Geneious v8.1.5 (Biomatters, Auckland, New Zealand). The consensus sequences were compared against the NCBI database (www.ncbi.nlm.nih.gov) using standard nucleotide BLAST (Altschul et al., 1990) to confirm sequence similarity and adenovirus identity. Specific parameters used for BLAST searches is as follows; optimization-somewhat similar sequences (blastn), E-value threshold-0.05. All other general, filtering and masking parameters were kept as default.

### 2.3 Sequence analyses: phylogenetic reconstruction and haplotype network

We further investigated the evolutionary relationship between the sequences obtained from this study and other known adenovirus sequences. A 242 bp short fragment DPOL gene alignment was created using the MAFFT alignment tool (v7.490) (Katoh & Standley, 2013) after being cleaned using Gblocks online tool with default settings (Talavera & Castresana, 2007) to detect and remove poorly aligned regions (e.g., gaps). In addition to the Murine adenoviruses, other known mammalian adenovirus sequences (Mastadenoviruses) and one Fowl adenovirus (Aviadenovirus; KT862812) sequence were downloaded from NCBI and included in the alignment. The best nucleotide substitution model for the alignment was determined, and phylogenies were reconstructed using IQ-TREE v1.6.12 (Trifinopoulos et al., 2016) with 1000 bootstrap replicates. The resulting bootstrap consensus tree was visualized and annotated using ITOL (Letunic & Bork, 2021). The Fowl adenovirus was used as an outgroup to root the phylogenetic tree.

Besides, we reconstructed a median-joining haplotype network to understand the relationship between sequences generated in this study. The alignment of DPOL gene (242 bp) was imported to POPART (Leigh & Bryant, 2015), and a median-joining network was created.

### 2.3. Pairwise genetic distance and private mutations

We calculated pairwise genetic distance (p-distance) between the sequences generated in this study and other murine adenoviruses (NC_012584 & NC_014899). We used a maximum composite likelihood model with uniform substitution rates and 1000 bootstrap iteration in MEGA 11 (Tamura et al., 2021). Using FastaChar (Merckelbach & Borges, 2020), we further identified private mutations in the short fragment of DPOL gene within our samples in comparison with other murine adenoviruses

## 3. Results

### 3.1 Adenovirus prevalence in small mammals

We detected the presence of adenovirus only in pooled tissue samples, all swabs tested negative. Adenovirus prevalence varied among small mammals tested (0-38.8%, Table 1) with an overall prevalence of 21.21% (n=132). Prevalence was calculated as the proportion of individuals positive for adenovirus in the samples tested. Interestingly, we observed a high prevalence of adenovirus in *R. satarae* (38.8%, n= 66), an endemic small mammal in the Western Ghats, as compared to other species in the community (Table 1). Among the other species in the community, *Mus cf. terricolor* showed high prevalence (16.6%, n=6) followed by *Mus cf. fernandoni* (4.3%, n=22). None of the *R. rattus* and *S. niger* tissue samples tested were positive for adenovirus, potentially due to lower prevalence and low sample size.

**Table 1:** Adenovirus prevalence in various small mammals tested from Kadamane forest-plantation mosaic. Pooled tissue consists of liver, spleen, kidney, lungs, and intestine samples.

|  | Adenovirus prevalence: number of positives / number of samples tested (%) |  |  |
| --- | --- | --- | --- |
|  | Pooled tissue | Oral swabs | Rectal swabs |
| *Rattus satarae* | 26/66 (38.8) | 0/64 (0) | 0/66 (0) |
| *Mus cf. fernandoni* | 1/22 (4.3) | 0/2 (0) | 0/2 (0) |
| *Mus cf. famulus* | 0/26 (0) | 0/2 (0) | 0/6 (0) |
| *Mus cf. terricolor* | 1/6 (16.6) | - | - |
| *Rattus rattus* | 0/10 (0) | - | 0/8 (0) |
| *Suncus niger* | 0/2 (0) | - | 0/2 (0) |
| Total | 28/132 (21.2) | 0/68 | 0/84 |

#### Phylogenetic relationships, genetic distance and private mutations

All the 28 positive samples identified (26 from *R. satarae*, one from *M. cf. fernandoni*, and one from *M. cf. terricolor*) were verified for adenovirus using BLAST search (Table S1), The BLAST results suggested 75-78% similarity with Murine adenoviruses sequences (E value: 7e^-^55), classified as a part of the Mastadenovirus group by the International Committee on Taxonomy of Viruses (ICTV). However, only the 26 samples generated from *R. satarae* showed satisfactory quality (>90%) for phylogenetic analysis. All these sequences are deposited in GenBank (www.ncbi.nlm.nih.gov) under the accession number OR906154 - OR906179.

Our phylogenetic analysis revealed five distinct adenovirus lineages from *R. satarae* (Figure 2A) belonging to six haplotypes (Figure 2B). Collectively, all these lineages share common ancestry with Murine adenovirus (NC_012584) in our phylogeny. Lineage 1 showed phylogenetic affinity with Murine adenovirus (NC_012584) by forming a sister taxon (p-distance = 0.27, S2). The remaining lineages (2-5) formed a distinct monophyletic group, with lineages four and five being sister to each other, while lineages two and three are basal to this cluster (Figure 2A). All these four lineages had significant genetic distance (p distance) of 0.33 - 0.37 (mean = 0.355) from Murine adenovirus (NC_012584; Table S2).

**Figure 2:**
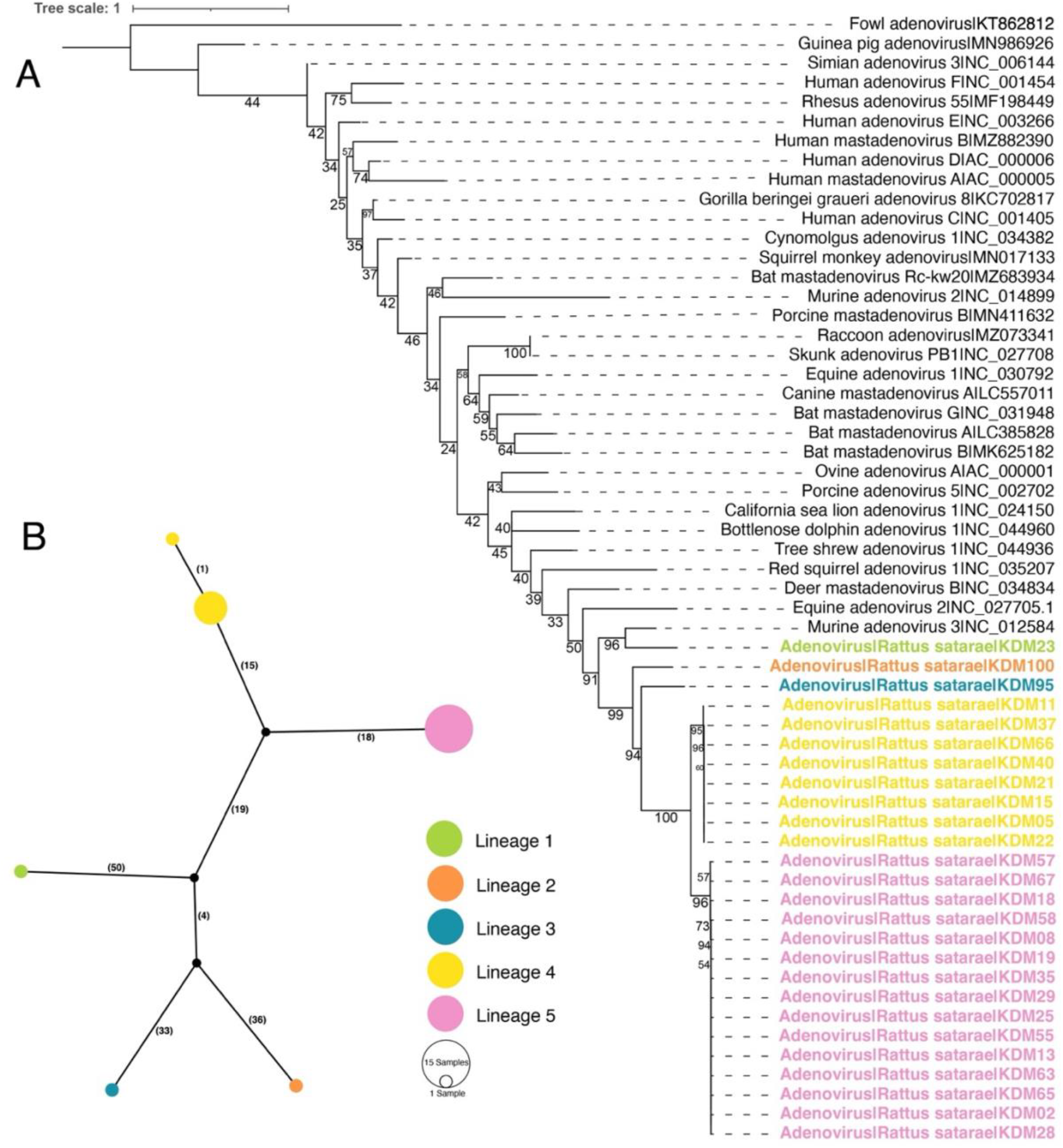
(A) Maximum likelihood phylogenetic tree of adenovirus based on 242 bp of DPOL short fragment. The phylogeny was rooted using Fowl adenovirus (KT862812). Numbers around nodes represent bootstrap values. Unique lineages identified in this study are colored differently. (B) A haplotype network showing different adenovirus lineages (DPOL) identified in *R. satarae*. The number next to the edges indicates the number of nucleotide differences between haplotypes. Colors in the haplotype network correspond to the novel lineages in the phylogenetic tree identified in this study.

Most of the adenovirus sequences from *R. satarae* (n=23) were part of lineage four and five represented by three unique haplotypes (Figure 2B). The remaining three sequences formed three unique haplotypes (p-distance = 0.3-0.38, mean = 0.34) corresponding to lineage one, two, and three. Further, we identified six to 17 private mutation which are unique to the new lineages described in this study (Table S3).

## 4. Discussion

In this study, we used molecular detection and sequencing to characterize circulating adenovirus diversity in small mammals inhabiting human managed forest-plantation mosaic in southern India. Our results reveal previously undetected adenovirus diversity associated with endemic small mammals inhabiting the Western Ghats biodiversity hotspot.

We report a high overall prevalence of adenovirus in the small mammal community with high prevalence in the most abundant small mammal *Rattus satarae*, potentially suggesting density-dependent transmission as observed in numerous other viral and host systems (Renwick et al., 2007). Interestingly, within our datasets, all individuals of *R. rattus*, a synanthropic species known to carry several virulent pathogens (Gravinatti et al., 2020), tested negative for adenoviruses, aligning with the previously reported *Bartonella* prevalence in the species (Ansil et al., 2021).

Till date, three prominent adenoviruses from small mammals have been characterized (*Murine adenovirus 1, Murine adenovirus 2*, and *Murine adenovirus*) globally with the advent of whole genome sequencing (Hemmi & Spindler, 2019). Recent reports suggest a widespread diversity of adenoviruses associated with small mammals, especially from tropical areas (Diffo et al., 2019; Ochola et al., 2022; Zheng et al., 2016), through sequencing of a short fragment of the DNA polymerase gene. The phylogenetic pattern observed in our dataset supported the existence of five distinct adenovirus lineages in *R. satarae*, all of which showed genetic differences (more than 25%) substantially higher than the cutoff of 5% -15% difference in polymerase gene to be considered as distinct lineages (Diffo et al., 2019). The presence of private mutations within these lineages (Table S3), along with their unique phylogenetic position (monophyletic lineages, except lineage one), further substantiate our results.

Given these results, we strongly recommend the isolation and further characterization of adenoviruses from *R. satarae* from the Western Ghats. High prevalence of adenoviruses in small mammals in these mixed-use landscapes have major implications for the health of other wildlife including small carnivores (secondary consumers) which inhabit these landscapes, as they can acquire infections through feeding on infected hosts (Thiry et al., 2007). We contend that long-term efforts with broader spatial coverage are essential to elucidate the role of species diversity, as opposed to species identity, in shaping the dynamics and evolution of novel adenovirus variants in mixed-use landscapes. Our study is one of the few initial attempts to understand the adenovirus diversity in wildlife in the region, which can provide impetus to establish standardized model systems to investigate the implications of the novel adenovirus variants.

## Acknowledgments

We also acknowledge the support of NCBS Sanger sequencing facility. ABR was supported by the SPM fellowship by CSIR-HRDG, Government of India. AS was supported by fellowship from the Trivedi School of Biosciences, Ashoka University. KMG acknowledges the support of DBT-Ramalingaswami Fellowship (BT/HRD/35/02/2006).

## Data availability statement

Sequence data generated in this study is available under GenBank accession number OR906154 - OR906179

