## Supplementary Tables for "Identification of distinct rodent-associated adenovirus lineages from a mixed-use landscape in the Western Ghats biodiversity hotspot"

**Table S1**: Top 5 blast search results for representative sequences belonging to unique haplotypes

| | Scientific name (as per NCBI) | Query cover | E value | Percentage identity | Accession |
|---|---|---|---|---|---|
| Haplotype 1 (Lineage 1) OR906164 | Adenovirus PREDICT_AdV-25 | 88% | $8e^{-34}$ | 75.64% | MK325864 |
| | *Murine adenovirus 3* | 96% | $1e^{-30}$ | 72.66% | OR405320 |
| | *Murine adenovirus 3* | 96% | $1e^{-30}$ | 72.66% | NC_012584 |
| | *Murine adenovirus 3* | 96% | $1e^{-30}$ | 72.66% | EU835513 |
| | *Mastadenovirus sp* | 82% | $2e^{-28}$ | 74.89% | MK249326 |
| Haplotype 2 (Lineage 2) OR906179 | Adenovirus PREDICT_AdV-46 | 92% | $2e^{-21}$ | 70.49% | MK325905 |
| | *Mastadenovirus sp* | 92% | $2e^{-21}$ | 70.49% | MK249337 |
| | *Bat mastadenovirus* | 60% | $1e^{-13}$ | 73.29% | MN490088 |
| | *Mastadenovirus sp* | 72% | $1e^{-13}$ | 70.47% | MK249325 |
| | Adenovirus PREDICT_AdV-21 | 71% | $4e^{-12}$ | 70.00% | MK325863 |
| Haplotype 3 (Lineage 3) OR906178 | Adenovirus PREDICT_AdV-44 | 99% | $2e^{-35}$ | 73.80% | MK325798 |
| | Adenovirus PREDICT_AdV-44 | 99% | $7e^{-35}$ | 73.70% | MK325797 |
| | Adenovirus PREDICT_AdV-44 | 99% | $1e^{-32}$ | 73.06% | MK325794 |
| | *Mastadenovirus sp* | 92% | $5e^{-30}$ | 73.12% | MK249316 |
| | *Mastadenovirus sp* | 92% | $5e^{-30}$ | 73.12% | MK249315 |
| Haplotype 4 (Lineage 4) OR906170 | Adenovirus PREDICT_AdV-44 | 100% | $6e^{-35}$ | 73.95% | MK325798 |
| | Adenovirus PREDICT_AdV-44 | 100% | $6e^{-35}$ | 73.95% | MK325797 |
| | Adenovirus PREDICT_AdV-44 | 100% | $3e^{-33}$ | 73.56% | MK325794 |
| | Adenovirus PREDICT_AdV-46 | 94% | $5e^{-30}$ | 73.77% | MK325905 |
| | *Mastadenovirus sp* | 94% | $5e^{-30}$ | 73.77% | MK249337 |
| Haplotype 5 (Lineage 4) OR906157 | Adenovirus PREDICT_AdV-44 | 99% | $2e^{-34}$ | 73.85% | MK325794 |
| | Adenovirus PREDICT_AdV-44 | 99% | $9e^{-33}$ | 73.46% | MK325798 |
| | Adenovirus PREDICT_AdV-44 | 99% | $9e^{-33}$ | 73.46% | MK325797 |
| | *Mastadenovirus sp* | 94% | $6e^{-29}$ | 72.98% | MK249312 |
| | Adenovirus PREDICT_AdV-46 | 93% | $2e^{-28}$ | 73.25% | MK325905 |
| Haplotype 6 (Lineage 5) OR906171 | Adenovirus PREDICT_AdV-46 | 90% | $5e^{-30}$ | 73.58% | MK325905 |
| | *Mastadenovirus sp* | 90% | $5e^{-30}$ | 73.58% | MK249337 |
| | Adenovirus PREDICT_AdV-44 | 99% | $2e^{-29}$ | 72.59% | MK325794 |

| | | | | |
|---|---|---|---|---|
| Adenovirus PREDICT_AdV-44 | 99% | $8e^{-28}$ | 72.22% | MK325798 |
| Adenovirus PREDICT_AdV-44 | 99% | $8e^{-28}$ | 72.22% | MK325797 |

**Table S2**: Genetic distance of (p-distance) between representative sequences from each lineage and two known murine adenoviruses. Note that, Murine adenovirus NC_014899 and NC_012584 are not monophyletic in our phylogenetic analysis (Figure 2A).

| | Murine adenovirus (NC_014899) | Murine adenovirus (NC_012584) | Lineage 1 (OR906164) | Lineage 2 (OR906179) | Lineage 3 (OR906178) | Lineage 4 (OR906170) | Lineage 5 (OR906171) |
|---|---|---|---|---|---|---|---|
| Murine adenovirus (NC_014899) | | | | | | | |
| Murine adenovirus (NC_012584) | 0.49 | | | | | | |
| Lineage 1 (OR906164) | 0.43 | 0.28 | | | | | |
| Lineage 2 (OR906179) | 0.51 | 0.36 | 0.36 | | | | |
| Lineage 3 (OR906178) | 0.53 | 0.34 | 0.38 | 0.30 | | | |
| Lineage 4 (OR906170) | 0.44 | 0.37 | 0.37 | 0.33 | 0.31 | | |
| Lineage 5 (OR906171) | 0.43 | 0.37 | 0.37 | 0.31 | 0.33 | 0.16 | |

**Table S3:** Private mutations identified in the Adenovirus isolated from *Rattus satarae* samples in comparison with the known Murine Adenoviruses (GenBank accession number NC_012584 and NC_014899)

| | Number of private mutations | Details of private mutation | |
|---|---|---|---|
| | | Location | Mutation |
| **Lineage 1** | 17 | 21 | C |
| | | 39 | G |
| | | 54 | C |
| | | 55 | G |
| | | 56 | C |
| | | 67 | G |
| | | 69 | G |
| | | 75 | C |
| | | 77 | G |
| | | 79 | T |
| | | 80 | C |
| | | 127 | G |
| | | 147 | G |
| | | 150 | G |
| | | 155 | C |
| | | 239 | T |
| | | 240 | T |
| **Lineage 2** | 17 | 10 | T |
| | | 15 | T |
| | | 22 | A |
| | | 46 | A |
| | | 47 | T |
| | | 57 | A |
| | | 66 | C |
| | | 75 | A |
| | | 76 | A |
| | | 80 | T |
| | | 90 | G |
| | | 112 | T |
| | | 147 | A |
| | | 157 | A |
| | | 162 | T |
| | | 166 | A |
| | | 204 | G |
| **Lineage 3** | 17 | 15 | G |
| | | 62 | C |
| | | 67 | T |
| | | 68 | T |
| | | 69 | T |
| | | 73 | A |
| | | 81 | A |
| | | 90 | T |
| | | 102 | T |
| | | 106 | T |
| | | 107 | C |
| | | 112 | A |
| | | 113 | A |
| | | 156 | T |
| | | 213 | T |
| | | 222 | A |
| | | 237 | G |

| | | | |
|---|---|---|---|
| **Lineage 4** | 7 | 42 | A |
| | | 78 | T |
| | | 81 | T |
| | | 99 | C |
| | | 105 | A |
| | | 174 | C |
| | | 189 | T |
| **Lineage 5** | 6 | 21 | A |
| | | 51 | G |
| | | 84 | C |
| | | 189 | C |
| | | 219 | G |
| | | 240 | G |
